# Environmental DNA from a small sample of reservoir water can tell volumes about its biodiversity

**DOI:** 10.1101/2021.08.06.455478

**Authors:** André lq Torres, Danielle Las Do Amaral, Murilo Guimaraes, Henrique B Pinheiro, Camila M Pereira, Giovanni M De Castro, Luana Ta Guerreiro, Juliana A Americo, Mauro F Rebelo

## Abstract

We evaluated the potential of metabarcoding in assessing the environmental DNA (eDNA) biodiversity profile in the water column of an hydroelectric power plant reservoir in southeast Brazil. Samples were obtained in three technical replicates at 1 km from the dam at 1, 13 and 25 m depths. For each minibarcodes -- COI, 12S and 16S -- 1.5 million paired-reads (150 base pairs) were sequenced. A total of 44 unique taxa were found. COI identified most of the taxa (34 taxa; 77.2 %) followed by 16S (14; 31.8 %) and 12S (10; 22.7 %). All minibarcodes identified fishes (13 taxa), however, COI detected other aquatic macro-invertebrates (18), algae (3) and amoebas (2). Richness was the same across the three depths (35 taxa), although, beta diversity suggested slightly divergent profiles. In just one location we identified 15 taxa never reported previously, 50% of the fish species identified in the last year of fishery monitoring and 13% of the species in biodiversity surveys performed from 2012 to 2021. Clustering into Amplicon Sequence Variants (ASV) showed that 12S and 16S are able to detect predominant haplotypes of fishes, suggesting they are suitable to study population genetics of this group. In this study we reviewed the species occurring within the Três Irmãos reservoir according to previous conventional surveys and demonstrated that eDNA metabarcoding can be applied to monitor its biodiversity.

## 1. INTRODUCTION

Hydroelectric power plant (HPP) reservoirs require constant monitoring for operational and environmental purposes, which poses challenges for operators and regulators [1,2]. The so-called Scientific fisheries for morphological identification require euthanization of fishes and are quite invasive [3]. Such collection techniques are unlikely to capture the true species richness due to intrinsic biases [4] as gillnets often underrepresent large adults and small fishes [3,5]. Morphological identification relies upon the availability and knowledge of taxonomists; it can be a laborious, slow and expensive process, with high error rates [6].

Environmental DNA (eDNA) is the shed DNA molecules in the water, soil or air originating from mucus, feces, skin peeling or other biological sources [7]. The use of eDNA as a source of biological material to identify biodiversity, including fishes, in aquatic environments has been growing due to its lower cost and invasiveness [5,8]. When the number of taxa identified from environmental DNA is compared to the consolidated result of multiple traditional capture methods, one finds that as little as 1 L of water can contain DNA representative of 85% of the fish biodiversity in lakes [9].

Indeed, environmental DNA can detect comparable or greater species richness [3,10], including rare species [11]. These DNA databases are enormous and growing: the Barcode of Life Data System (BOLD) has 324,988 species with barcodes, including 22,986 bony fish species [12,13]. Studies have shown that eDNA can increase species identification, reducing error due to cryptic species or juveniles and eDNA can better capture fish biodiversity in hydroelectric impoundments [3,14].

The aim of the study was to start an inventory of the biodiversity of the Três Irmãos (TI) reservoir using eDNA as a source of biological material. The results demonstrated that eDNA metabarcoding has potential to survey aquatic biodiversity of lentic environments such as HPP reservoirs. The results shed light on key issues that will guide the larger scale application of this methodology at the reservoir, such as the molecular marker taxa coverage, number of replicates and depths to be sampled for the optimization of the biodiversity inventory.

### 2. MATERIAL AND METHODS

### 2.1. The reservoir

Três Irmãos is the largest of the 6 reservoirs built in the Tietê river, São Paulo State, southeast Brazil (Figure 1), with a flood area of 817 km^2^, a volume of 13,800 x 10^6^ m^3^ and an average depth of 17 m [15]. The fish biodiversity in the reservoir is constantly monitored for operational and management purposes, with the traditional sampling stations marked in the figure with numbers 1 to 4.

**Figure 1.**
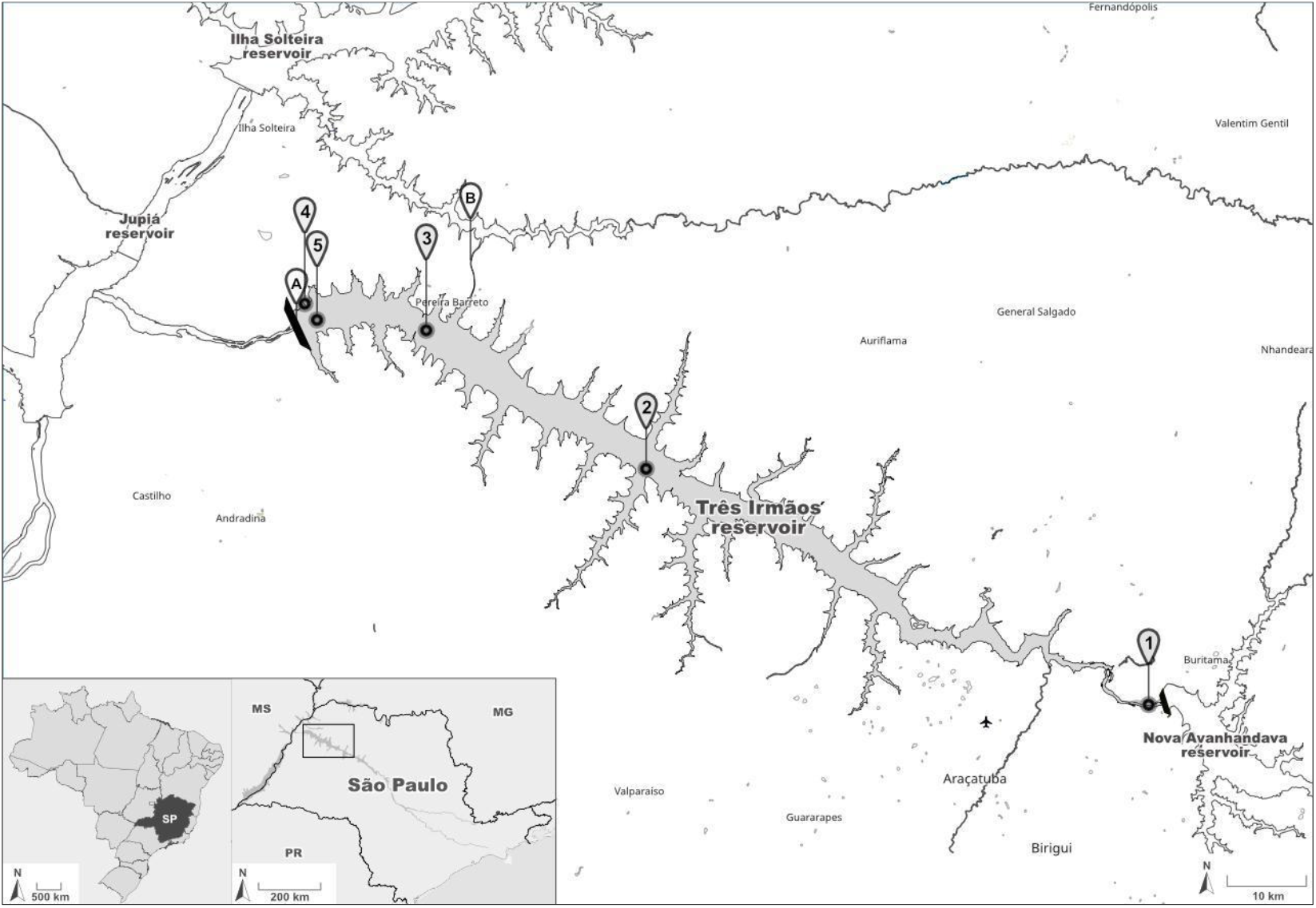
Três Irmãos (TI) reservoir in southeast Brazil (highlighted in gray). Point A shows the TI dam and (B) Pereira Barreto service channel with nearby Ilha Solteira reservoir. Points numbered 1 to 4 (1-P1-JNA; 2-P2-JAC; 3-P3-JBA; 4-P5-LIE) show where the traditional surveys were carried out by the hydroelectric company. Point 5 indicates the sampling location of this study. Small inset maps in the lower left corner show where the State of São Paulo is situated within Brazil and where the reservoir is located within São Paulo. The course of the Tietê River is shown and the TI reservoir is enclosed by a rectangle.

The eDNA sampling point was located approximately 1 km from the center of Três Irmãos dam (20°40.077’S, 051°16.702’W; Figure 1). One liter of water was collected in 3 technical replicates at three depths: 1 m, 13 m and 25 m from the surface, utilizing a Van Dorn bottle (Nalgon, Cat. No. 2330). Water samples were promptly filtered through 0.22 µm Sterivex filters (Merck-Millipore, Cat.no. SVGP01050) in 60 mL increments with disposable syringes. After filtration, remaining liquid was expelled by injecting air and the filter cartridges were then filled with sterile Longmire’s buffer [16] and kept at 4°C until DNA extraction.

### 2.2. Biodiversity Inventory of the Três Irmãos Reservoir

In an effort to have a reference baseline of the known biodiversity of the Três Irmãos reservoir, we conducted a survey of species that have been previously documented in this region by means of other traditional methods. Two strategies were used: 1) a search, limited to the geographic area of the reservoir, in the Global Biodiversity Information Facility (GBIF) database [17], and 2) a non-systematic literature review, which involved the search and review of articles, theses, and reports on monitoring of ichthyofauna and fisheries, the latter provided by hydroelectric concessionaires. The GBIF search was done by selecting an area of the map that encompasses the reservoir and a small region beyond (downstream from) the dam, as some species may traverse the dam and be reported there. The area can be seen in https://doi.org/10.15468/dl.g58exn.

### 2.3. Environmental DNA extraction and Deep sequencing

DNA extraction from the Sterivex filters were carried out using the DNeasy® Blood & Tissue kit (Qiagen) with adaptations previously described in the Environmental DNA Sampling and Experiment Manual (Version 2.1) [18]. DNA concentration was determined using Qubit dsDNA HS Assay Kit (Thermo Fisher Scientific). The amplicons from mitochondrial genes targeting the minibarcodes of cytochrome *c* oxidase I (COI), 16S rRNA gene, and 12S rRNA gene were sequenced. The sequences of the primers used for metabarcoding are described in supplementary table 2. Amplicons were indexed using Nextera XT indexes (IDT for Illumina). Library quality control was performed using Tapestation (Agilent). DNA concentration was normalized and all amplicons were pooled in one library. The library was sequenced on Illumina NovaSeq S4 Flow Cell at the 2X150 configuration.

### 2.4. Bioinformatic analysis

All fastq files of each minibarcode were imported into the QIIME2 pipeline [19] with their respective depth, barcode and sample date associated through the metadata file. The forward and reverse primers were trimmed from the reads using the DADA2 [20]. It was also used to denoise reads, merge the paired ones into amplicons and remove chimeras. The resulting amplicon dataset was submitted to the Amplicon Sequence Variant (ASV) clustering, which was also carried out with DADA2. The taxonomy identification of the ASVs was performed using three different methods: 1) Naive Bayesian classifier [21] trained with sequences dataset of the 12S Mitohelper [22], COI classifier version 4.0 [23], and the MIDORI 16S dataset (release 242; [24]); 2) BLASTn searches [25] against the curated mitochondrial reference database available in NCBI (https://ftp.ncbi.nlm.nih.gov/refseq/release/mitochondrion/ [26]; and the nucleotide non-redundant (nt) database [27]. Only hits with e-value lower than 1e-50 and query cover higher than 90% were allowed. ASVs with no match, off-target and non-eukaryotic sequences were removed from the dataset; 3) Phylogenetic analysis was carried out with the representative sequences of the ASVs of fish using the minibarcode sequence extracted from mitochondrial genomes and sequences from voucher species as reference. Sequences were aligned with MAFFT [28]. All trees were built with the neighbor joining method [29] using the evolutive model kimura 2 parameters (K2P; [30]). Branch support was calculated performing 10,000 replicates of bootstrap. *Homo sapiens* was used as an outgroup to root the trees. Tree visualization and ASV frequency comparison was carried out with iTOL [31].

### 2.5. Statistical analysis and Data visualization

Taxonomic tree of all species found with the ASV clustering was built using PhyloT2 (https://phylot.biobyte.de). It was loaded into the iTOL together with the dataset containing the normalized Log10 (frequency sum) of the ASVs frequencies of each taxa found by the three minibarcodes. The rarefaction curves were calculated using the alpha-rarefaction plugin available in the QIIME2 pipeline [19] allowing up to 4,000 amplicons sampling to the 12S minibarcode, 560,000 to the 16S and 500,000 to the COI. The unrooted phylogenetic tree of each minibarcode was provided to calculate the Faith’s phylogenetic diversity. Concerning biodiversity, the number of species among the three communities found within the three depths was compared using a Poisson GLM. Alpha diversity Shannon Index (H’
s), beta diversity (Jaccard similarity) and the Pielou evenness were computed. The effect of depth on communities was estimated using Permutational Multivariate Analysis of Variance. The diversity metrics were obtained using vegan package [32] with a presence/absence matrix of all taxa found. Especifically for fishes, the probability of finding species in relation to depth and number of samples was estimated using a Binomial GLM. All biodiversity analyses were performed in R [33].

## 3. RESULTS AND DISCUSSION

Sampling at a single collection site in the reservoir, eDNA metabarcoding was capable of detecting 50% (n=8) of the fish species surveyed in 2020 (n=16) by the hydroelectric concessionary with traditional capture methods in 3 monitoring campaigns and 13% of the species found (n=59) over a ten year period (2012-2021) [34].

The eDNA sequencing results are presented, as well as how the bioinformatics pipeline was used to attribute the sequence reads to molecular markers that enabled taxa identification. Next, biodiversity profiles are shown for each sample, discussing richness and diversity indexes, and how they contribute to the biodiversity inventory of the reservoir. The potential of the environmental DNA approach, its limitations, and the likelihood of false positives and false negatives are discussed. The fish inventory, the group that is more relevant for biodiversity management of the reservoir, is more detailed and the potential to identify intraspecific variation in this group is also presented.

### 3.1 Sequencing and bioinformatics

Deep sequencing yielded an average of 1.5 million of 150 bp paired-end reads per sample obtained at 1, 13 and 25 m of depth near the TI dam. The read cleaning, amplicon merging and chimera removal steps yielded approximately 219,000 12S amplicons (1.4%), 7 million COI (47.4%) and 8.4 million 16S amplicons (61.3%) (supplementary table 3). In spite of the low throughput, valid 12S amplicons were obtained and further analysed. Clustering of identical amplicons resulted in 615 Amplicon Sequence Variants (ASVs) of the 12S, 4,404 ASVs of the COI and 2,993 ASVs of the 16S amplicons. After identifying the taxons with Bayesian classifiers and BLASTn searches, we performed a manual inspection and removed off-target and non-eukaryotic ASVs, yielding 62 (10.8%), 187 (4.2%) and 832 (27.7%) valid ASVs to the 12S, COI, and 16S minibarcodes, respectively (supplementary table 4). The merging of ASVs assigned to the same taxonomic group resulted in a total of 44 unique taxa, of which 10 (22.7%) were identified by the minibarcode 12S, 34 (77.2%) by the COI and 14 (31.8%) by the 16S. Amplicon length, nucleotide sequence and taxonomy identification are available in supplementary table 4.

### 3.2 Biodiversity profiles

The phylogenetic tree in Figure 2 shows the richness and abundance of DNA barcodes of each of the 44 taxa identified in each of the 9 sequenced samples. It represents the first overall eDNA biodiversity profile of the reservoir. Twenty-four taxa were classified at the species level, including 7 mammals, 1 avian, 8 fishes, 4 crustaceans, 3 rotifers and 1 annelid (Figure 2A); 6 at the genus level (4 fishes, 1 crustacean and 1 rotifer), and 2 at family level (Clupeidae and Naididae). Twelve were assigned to higher levels: 3 to order (Hemiptera, Neuroptera and Ploimida), 3 to class (Ostracoda, Bangiophyceae and Florideophyceae), 1 to subclade (Copepoda), 1 to clade (stramenopiles), and 3 to phylum (Chlorophyta, Tubulinea and Discosea), as shown in Figure 2A. The frequency of DNA fragments in the heatmap represents a rough estimation of the abundance of the species, even though this DNA abundance should be interpreted with caution as it may reflect body size of individual species [35] or drift [36]. Taxa richness was similar at each of the 3 depths (Poisson Generalized Linear Model β_depth_ = -4.334e-17, z = 0, p = 1),, with each community presenting 35 taxa and according to the. Alpha diversity (Shannon index H’) was similar across the three depths with values 3.48 for 1 m, 3.49 for 13 m and 3.45 for 25 m. Beta diversity (Jaccard similarity index) was 0.79 (p = 0.01) between 1 m and 13 m depth, 0.70 (p = 0.30) between 13 m and 25 m and 0.59 (p = 0.13) between 1 m and 25 m. The Pielou species evenness was 0.98 for both 1 m and 13 m, and 0.97 for 25 m. Overall, depth did not influence clustering within communities (PERMANOVA; F=0.12, df=2, p=0.89).

**Figure 2.**
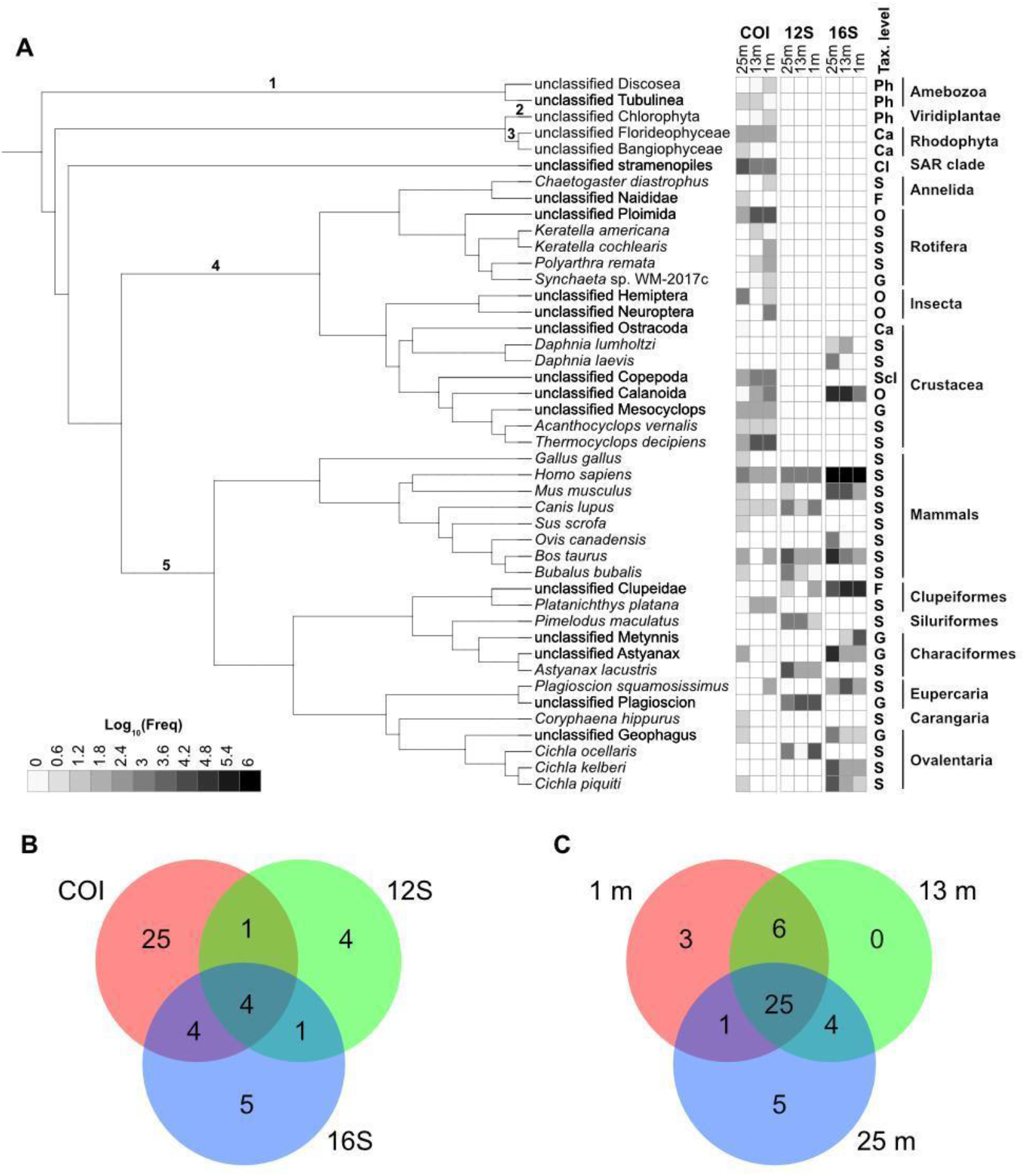
Taxonomy and species comparison per minibarcode by depth. A - Taxonomy of the organisms and biodiversity profile found at 25, 13, and 1 m depths for three minibarcodes (COI, 12S and 16S). The frequency -- actually sum of the frequencies in three replicates at each depth -- of each taxon by molecular marker is shown using a[Log_10_ grey scale. Taxonomy levels are represented by Ph (phylum), Ca (class), Scl (subclass), Cl (clade), O (order), F (family), G (genus) and S (species). Solid black lines highlight the taxonomic groups. Numbers in the internal nodes of the tree indicate the Amoebozoa clade (1), Viridiplantae kingdom (2), Rhodophyta phylum (3), protostomes (4) and deuterostomes (5) clades. B - Venn diagram shows the comparison of the species found by the three minibarcodes. C - Venn diagram shows the comparison of the species across the three depths.

The minibarcode Venn diagram (Figure 2B) shows that COI alone identified more taxa than any other minibarcode, a total of 25. Most are protostomes (arthropodes, rotifers and annelids) and COI was essential to assess the diversity of macroinvertebrates and plankton. Five species were found only by the 16S minibarcode, while 4 species, mostly fish and mammal groups, were exclusively found by the 12S minibarcode. Four species were identified by both 16S and COI, 1 by 12S and COI, and 1 by 12S and 16S. Only 4 species were detected by all minibarcodes (Figure 2B). These results stress the importance of using more than one marker to improve the species detection capacity, with 12S and 16S recommended as the most universal for detecting fish species [5].

The depth Venn diagram (Figure 2C) shows that most species, 25 in total, were identified at all 3 depths, with 3 being only at the surface (1 m) and 5 only at the bottom (25 m). It is tempting to look for an ecological explanation for the presence of some species in some of the samples, but while factors like species occupancy [37,38], eDNA persistence [39,40], quantity [5,41], and animal physiology [42] may influence the distribution pattern of eDNA, the absence of a species might also be the result of a primer bias [5], caused by failure to detect identical target DNA sequences in replicate samples, especially in samples with very low DNA concentrations [41]. Thus, while the presence of DNA in one sample is evidence of the existence of that species in that sample, the absence of a species in a given sample cannot rule out the existence of the species in it. The rarefaction curves (Supplementary figures 1 and 2) suggest that sequencing effort was sufficient to identify all species in each sample, and support the differences across the three depths.

Additionally, the probability of finding the group of fish species in water samples varied when considering one, two or three replicates and also within depths, between 0.28 and 0.41 (Figure 3). The highest probabilities of detecting the species were found in 25 m depth samples, with a 19.5% increase in the probability when considering three replicates of water sample, instead of only one.

**Figure 3.**
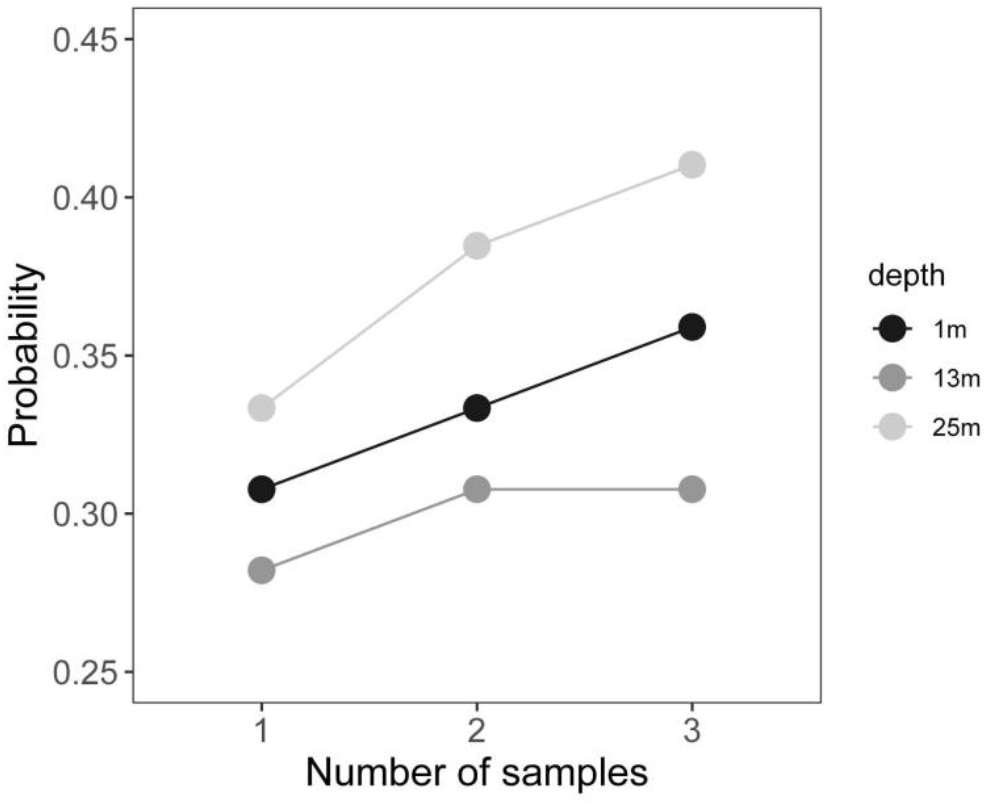
Probability of finding fish species per depth as a function of number of replicates.

The TI reservoir is a predominantly lentic environment, with water flow velocity lower than 0.05 m/s and little renovation of water as found by previous hydrodynamic modeling (unpublished data), leading to a tendency of silting, as previously observed [43]. eDNA behaves like fine particulate organic matter (FPOM) in regards to transport distance and deposition in riverbeds. Thus, the concentration of eDNA can be influenced by the hydrological profile of the water body as well as the breadth and velocity of water movement at different depths of the reservoir [44]. In general, the slower the velocity, the shorter the distance traveled before deposition [44]. Combined with the silting tendency, this may explain why we did not detect some species when sampling a single site.

Because eDNA metabarcoding permits the collection of more biological material per sample -- DNA both of more distinct organisms and more individuals of each -- than any other biological sampling method, and DNA sequencing and bioinformatics carries less analytical error than morphological identification, we would expect that, over time, the risk of false positive and false negatives will reduce. As non-invasive sampling with automated analysis can be performed more often, at a lower cost, the increase in the number of records should also increase the robustness of the conclusions derived from these observations.

### 3.3 Biodiversity Inventory

We constructed a list with 528 taxa previously reported in the literature and in the Global Biodiversity Information Facility (GBIF) database [17] that occur in the TI reservoir and the river basin (detailed in Materials and Methods). Then, we scrutinized GenBank and BOLD in order to identify the COI, 12S and 16S molecular markers for the species occurring in the TI reservoir (Supplementary table 1). Hereafter we used this list of species as a reference to discuss the detection or non-detection of taxa found by metabarcoding.

Fifteen taxa had their occurrence in the TI reservoir registered for the first time in this study. The olygochaeta Chaetogaster diastrophus, was registered in upstream reservoirs [45,46]. The rotifer Polyarthra remata was previously reported in other rivers of the Paraná basin [47] and Synchaeta sp. in Tietê headwaters [48] and upstream reservoirs (Nova Avanhandava and Promissão; [47]. Both endemic microcrustacean Daphnia laevis [49] and Mesocyclops sp. [50] have been reported in other upstream reservoirs. SAR (Stramenopila, Alveolata, and Rhizaria) organisms belonging to Stramenopila clade were not identified beyond this level, probably because the lack of reference sequences hampers a reliable taxonomic identification of this group [51].

A total of 21 taxa previously identified by morphological taxonomy were also identified by metabarcoding of eDNA (Table 1). Among the primary producers, we identified chlorophyta and rhodophyta (Bangiophyceae and Florideophyceae); both had already been reported in tributaries that flow into the Tietê river [52].

**Table 1:**
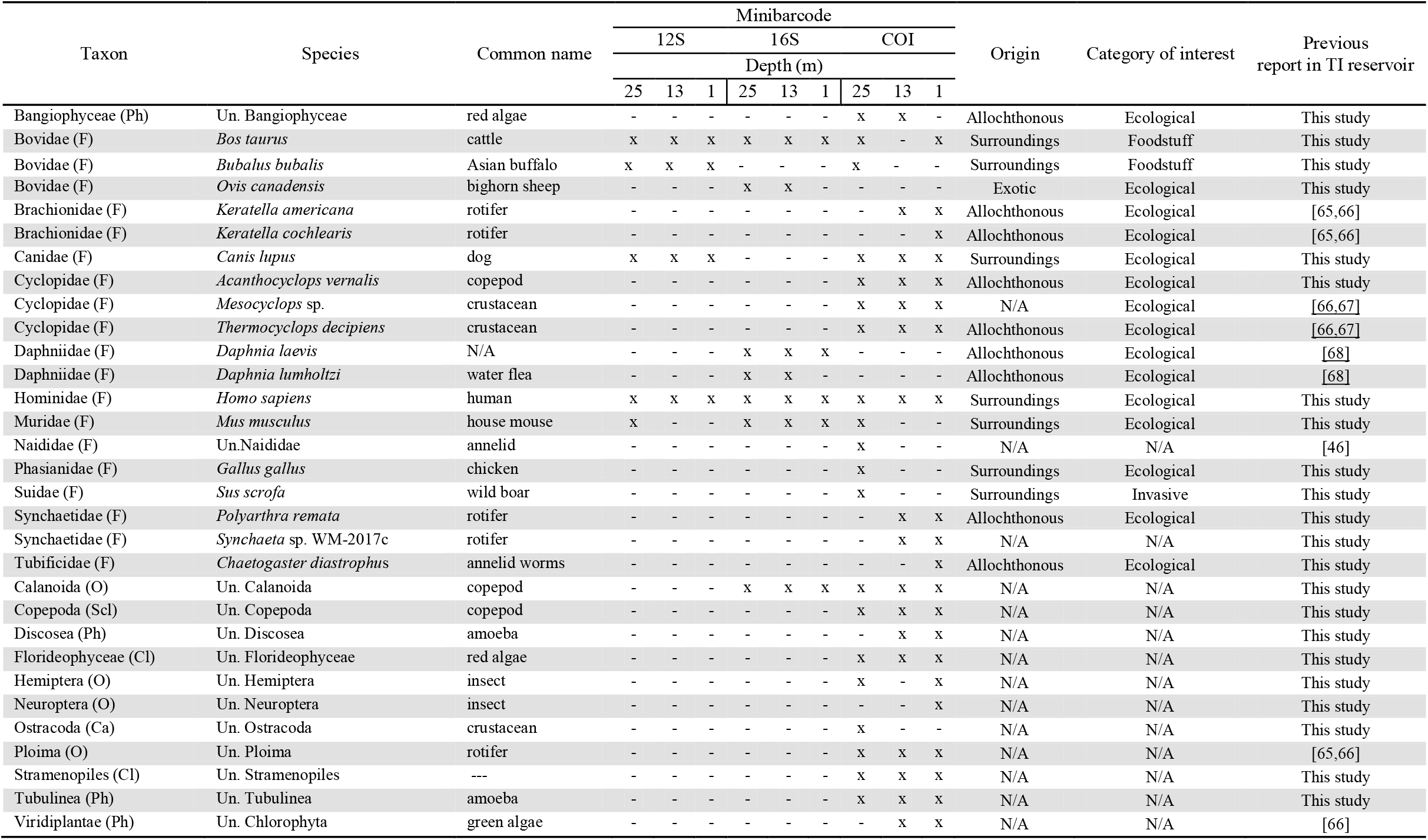
List of taxonomic groups identified in the water column of the TI reservoir through the eDNA and metabarcoding approach at 3 depths (25 m, 13 m and 1 m; m= meter). All organisms except fish are shown. Taxonomy levels are represented by Ph (phylum), Ca (class), Scl (subclass), Cl (clade), O (order), F (family), G (genus) and S (species), Un = unclassified, X=detected, - = non-detected, N/A = not applicable, due to impossibility to reach species or genus level of identification.

Among protostomes, we identified arthropods, rotifers and annelids, which are commonly found in the Tietê river. According to studies of stomach contents these protostomes constitute part of the diet of fish [53]. Ten arthropods were identified including planktonic crustaceans like the exotic Daphnia lumholtzi, which is native to Australia [54,55]; Thermocyclops decipiens [56]; and the cosmopolitan Acanthocyclops vernalis [57]. Other crustaceans that form the zooplankton community were identified at higher phylogenetic levels, such as: the Calanoida order, which includes two families occurring in the Tietê river, Diaptomidae and Pseudodiaptomidae [58], the Ostracoda class [53] and Copepoda subclass, which is mainly represented by the cyclopoids in this region [59].

Within the Insecta class, we were only able to identify unclassified hemipteran and neuropteran [53]. Annelids were identified at the family level and belong to the Naididae family, which have been previously found in TI [46] and two upstream reservoirs (Ponte Nova and Bariri; [45]). All rotifers belong to the Ploimida order, with two species (Keratella americana and K. cochlearis) previously reported within the TI reservoir [47].

Some species that were present or predominant in previous reports in the literature were not detected through DNA sequences obtained in this work, such as Cyanophyta, which was predominant in 2011 [52], Heterokontophyta, and some aquatic plants. No macro-crustacean were detected, including the exotic Macrobrachium amazonicum and Macrobrachium jelskii, both previously found in TI [60]. Both odonata and coleopteran, previously reported in this area [53,61], were not registered. Molluscs [62] and Platyhelminthes also have been reported in the TI reservoir and in the stomach contents of fish living in this region [53] but none of the three minibarcodes were able to detect eDNA eDNA of these groups. Considering that the reservoir is infested with the invasive golden mussel Limnoperna fortunei [63], responsible for filtrating huge amounts of water and dispersing gametes in the water column, we would expect at least this species to be detected. It is possible that sedentary behaviour of these organisms and the low water circulation restrain their eDNA locally [44], but it is more likely that primers used in this study were not able to amplify these groups. We also found DNA of terrestrial mammals, such as humans, dogs, cows, sheep, and chickens. Their presence has been consistently reported in eDNA of water samples [14,64] and has probably been carried into the reservoir from the surroundings by sewage and run-off.

#### 3.3.1 Fish inventory

Due to the extensive fishing and aquaculture activities in the reservoir, as well as a need for better information for fisheries management, we dedicated more effort to the identification of fish in the samples. Phylogenetic analysis performed for each minibarcode validated that all 13 analyzed fishes belong to the Actinopterygii class, divided in 6 orders and 7 families (Supplementary figures 3, 4, 5; 12S, COI, and 16S, respectively), according to the most recent bony fishes classification [69] (Table 2). eDNA in this single location managed to retrieve 13 fish taxa (n=13) previously reported by traditional taxonomy, over the last 30 years in different parts of the TI reservoir (Supplementary table 1; Figure 1).

**Table 2:** Fish taxa list determined by eDNA through 12S, 16S and COI minibarcodes at 3 depths (25 m, 13 m and 1 m; m = meter). Un. = unclassified, x = detected, - = non-detected, N/A = not applicable, because it was not possible to reach species or genus level of identification, Autoc. = Autochthonous, Alloc. = Allochthonous.

| Order | Family | Specie | Common name | Minibarcodes |  |  |  |  |  |  |  |  | Origin | Migratory | Habitat | Commercial interest | Previous report |
| --- | --- | --- | --- | --- | --- | --- | --- | --- | --- | --- | --- | --- | --- | --- | --- | --- | --- |
|  |  |  |  | 12S |  |  | 16S |  |  | COI |  |  |  |  |  |  |  |
|  |  |  |  | Depth (m) |  |  |  |  |  |  |  |  |  |  |  |  |  |
|  |  |  |  | 25 | 13 | 1 | 25 | 13 | 1 | 25 | 13 | 1 |  |  |  |  |  |
| Characiformes | Characidae | *Astyanax lacustris* | Tambuí | x | x | x | - | - | - | - | - | - | Autoc. | No | Benthopelagic | Fishing | [73] |
| Characiformes | Characidae | *Astyanax* sp. | Tambari | - | - | - | x | x | x | x | - | x | N/A | N/A | N/A | N/A | [77,78] |
| Cichliformes | Cichlidae | *Cichla piquiti* | tucunaré azul, blue peacock bass | - | - | - | x | x | x | x | - | - | Invasive | No | Benthopelagic | Fishing | [72,73,79] |
| Cichliformes | Cichlidae | *Cichla kelberi* | tucunaré-amarelo, Kelberi peacock bass | - | - | - | x | x | x | - | - | - | Invasive | No | Benthopelagic | Fishing | [72,73,79] |
| Cichliformes | Cichlidae | *Cichla ocellaris* | tucunaré, furiba, butterfly peacock bass | x | x | x | - | - | - | - | - | - | Invasive | No | Benthopelagic | Fishing | [61] |
| Cichliformes | Cichlidae | *Geophagus* sp. | - | - | - | - | - | - | - | x | - | - | Alloc. | No | Benthic | Fishing | [72,73] |
| Clupeiformes | Clupeidae | *Platanichthys platana* | sardine, river plate sprat | - | - | - | - | - | - | x | x | x | Autoc. | No | Pelagic | Fishing | [80] |
| Clupeiformes | Clupeidae | unclassified Clupeidae | - | x | - | x | - | - | - | - | - | - | N/A | N/A | N/A | N/A | [80] |
| Carangiformes | Coryphaenidae | *Coryphaena hippurus* | carangaria, common dolphinfish | - | - | - | - | - | - | x | - | x | Alloc. | Yes | Pelagic-neritic | Fishing | This study |
| Siluriformes | Pimelodidae | *Pimelodus maculatus* | mandi catfish | x | x | x | - | - | - | - | - | - | Autoc. | Yes | Benthopelagic | N/A | [72,73,78,79,81] |
| Perciformes | Sciaenidae | *Plagioscion squamosissimus* | corvina, south american silver croaker | - | - | - | x | x | x | - | - | x | Alloc. | No | Benthopelagic | Fishing | [72,73,77,78,79,81] |
| Perciformes | Sciaenidae | *Plagioscion* sp. | - | x | x | x | - | - | - | - | - | - | N/A | N/A | N/A | Fishing | [72,73,77,78,79,81] |
| Characiformes | Serrasalmidae | *Metynnis* sp. | pacu-prata | - | - | - | - | x | x | - | - | - | N/A | N/A | N/A | Fishing/ornamental | [72,73,78,81] |

The 8 fish species detected belong to 5 orders previously identified within the reservoir: Siluriforme, Cichliforme, Eupercaria, Clupeiforme, and Characiforme. The most represented order was Cichliforme (3 species) and the only native species identified was the Siluriforme *Pimelodus maculatus*. Other 6 species are allochthonous, such as the (Eupercaria) *Plagioscion squamosissimus* (Corvina or silver Croaker) that was identified and is one of the most abundant species reported in ichthyofauna monitoring reports of the lentic environmental areas of the TI reservoir. *Astyanax* spp (Characiforme), known as Lambari, are constant, omnivorous, sedentary, and non-migratory species that have been reported in the reservoir and in the Tietê river since the 17^th^ century [70]. *Astyanax lacustris* was detected by eDNA and, at least, five *Astyanax* sp. were previously reported in the TI reservoir. We also identified *Platanichthys platana* (Sardine) (Clupeiforme), which was detected at the three depths and 3 species of *Cichla* sp. (Cichliforme), which are native from other Brazilian basins [71]: *Cichla piquiti, C. kelberi*, and *C. ocellaris*.

None of the 6 native fish species which are being introduced annually into the TI reservoir by the hydroelectric concessionaire as part of its restocking program were detected: *Piaractus mesopotamicus* (Pacu-guaçu), *Prochilodus lineatus* (Curimbatá), *Brycon orbygnianus* (Piracanjuba), *Pseudoplatystoma corruscans* (Pintado), *Salminus brasiliensis* (Dourado), *Leporinus elongatus* (Piapara). Data from the ichthyofauna and fishery monitoring program performed either by hydroelectric concessionaires from 1992 to 2020 [34], reviewed by Marques (2019;[72]) or species inventories conducted for different purposes [70,73], using traditional capture methods at the sampling location (P3-JBA) closest to ours, showed that the target species used in restocking programs are rare, representing less than 2% of total fish caught in 2019 [34]. So it is possible that the low population density of the target species could lead to false-negative results. Counterintuitively, we identified the marine carangaria (*Coryphaena hippurus*). It has been previously reported that sewage can carry DNA of marine fish that are part of nearby populations’ diet [9,74].

Beyond taxa identification and presence/absence determination, eDNA has been growing as a powerful tool to investigate intraspecific variation, especially when dealing with low-densities or threatened species, when non-invasive methods are preferable [75]. Amplicon clustering into ASVs allowed us to detect one haplotype of each freshwater fish identified by the COI minibarcode at the species level (*P. platana, C. piquiti* and *P. squamosissimus*; Supplementary figure 4). Twelve haplotypes of the *C. piquiti* and *C. kelberi* were detected by the 16S minibarcode (Supplementary figure 5) while 12S detected 7 of the *A. lacustris* and 8 of the *C. ocellaris* (Supplementary figure 3). The greater number of haplotypes recovered by the rRNA minibarcodes seems to be associated with the higher molecular diversity of both in comparison with COI, which is more conserved [76]. Although multiple haplotypes have been identified, only one was detected in all depths and at frequencies higher than the others for *A. lacustris, C. piquiti* and *C. kelberi* (Supplementary table 4). Three haplotypes were found for *C. ocellaris*, all at high frequencies, nonetheless they were predominant at the bottom (Supplementary table 4). This result indicates that both minibarcodes, 12S and 16S (primers used: Supplementary table 2), can be used to assess the intraspecific diversity of the local fish community, although we must consider the loss of resolution for some closely related species, e.g. *Astyanax* spp., and the lack of a reference sequence as was observed for clupeiformes, for example.

## 4. CONCLUSIONS

Metabarconding of environmental DNA from a single collection point in the reservoir was capable of detecting 50% of the fish species identified in 2020 by the hydroelectric concessionary using traditional capture methods. We believe that increasing the number of collection points across a wider swath of the reservoir and conducting collections during multiple seasons would likely increase the number and completeness of fish taxa detected by eDNA metabarcoding.

Previous unreported taxa, mainly from the planktoninc community, such as copepods and rotifers, were detected. With eDNA it was possible to simultaneously detect fishes and copepods, taxa that previously required completely separate methods of traditional sampling.

The use of 3 molecular markers increased the power of detection, with more taxa identified by COI. Still, 12S and 16S were important to detect fish. For fishes, the probability of detection increased with the increment in the number of sampling replicates. Thus, until we establish a monitoring protocol, we recommend that all monitoring efforts in the reservoir use at least 3 markers and perform independent samples in triplicate.

## 5. SUPPORTING INFORMATION

**Supplementary table 1. Inventory of fish and non-fish species in Três Irmãos Reservoir from literature and GBIF Search**

**Supplementary table 2. Primer pairs used to amplify the target amplicons in the COI, 12S rRNA, and 16S rRNA genes from eDNA of the TI reservoir**. Ta = Annealing temperature used.

**Supplementary table 3: N of paired-reads (input), N and % of high quality paired-reads (filtered), N of denoised paired-reads, N and % of merged paired-reads and N and % of non-chimeric paired-reads**

**Supplementary table 4: 12S, COI and 16S ASV clustering, amplicon length, frequency by depth, and classification**.

**Supplementary figure 1. Alfa rarefaction per sample of the 12S (A), 16S (B) and COI (C) minibarcodes**. Shannon index and the sequencing depth are represented by the y and x axes, respectively.

**Supplementary figure 2. Alfa rarefaction by depth (1, 13 and 25 m) of the 12S (A), 16S (B) and COI (C) minibarcodes**. Shannon index and the sequencing depth are represented by the y and x axes, respectively.

**Supplementary figure 3. Neighbor-joining tree using the minibarcode 12S region**. Leaves containing the ASVs belonging to fishes and reference sequences are coloured in black and red, respectively. *Pimelodus* sp. (Siluriforme) is represented by the clade A, *Astyanax lacustris* (clade B), Unknown clupeidae (Clade C), *Cichla ocellaris* (Clade D) and *Plagioscion* sp. (Clade E). Frequencies of each ASV within the replicates sampled at a depth of 1, 13 and 25 meters are shown at the right side of the leaves.

**Supplementary figure 4. Neighbor-joining tree using the minibarcode COI region**. Leaves containing the ASVs belonging to fishes and reference sequences are colored in black and red, respectively. *Coryphaena hippurus* is represented by the Clade A, *Plagioscion squamosissimus* (Clade B), *Platanichthys platana* (Clade C), *Astyanax* sp. (Clade D), *Geophagus* sp. (Clade E) and *Cichla piquiti* (Clade F). Frequencies of each ASV within the replicates sampled at a depth of 1, 13 and 25 meters are shown at the right side of the leaves.

**Supplementary figure 5. Neighbor-joining tree using the minibarcode 16S region**. Leaves containing the ASVs belonging to fishes and reference species are coloured in black and red, respectively. *Plagioscion squamosissimus* is represented by the clade A, *Geophagus* sp. (Clade B), *Cichla piquiti* (Clade C), *Cichla kelberi* (Clade D), *Metynnis* sp. (Clade E) and *Astyanax* sp. (Clade F). Frequencies of each ASV within the replicates sampled at a depth of 1, 13 and 25 meters are shown at the right side of the leaves.

## 6. ACKNOWLEDGMENTS

This work was financed by Tijoá Energia through the Brazillian National Electric Energy Agency ANEEL RD program (Grant PD-09151-2001/2020). We thank Angélica Beccato, Environment and Land Manager at Tijoá Energia, for her support and collaboration in the development of this work.

## Notes

### Competing Interest Statement

The authors have declared no competing interest.

