## Supplementary figures for "Environmental DNA from a small sample of reservoir water can tell volumes about its biodiversity"

### 1. Rarefaction curves

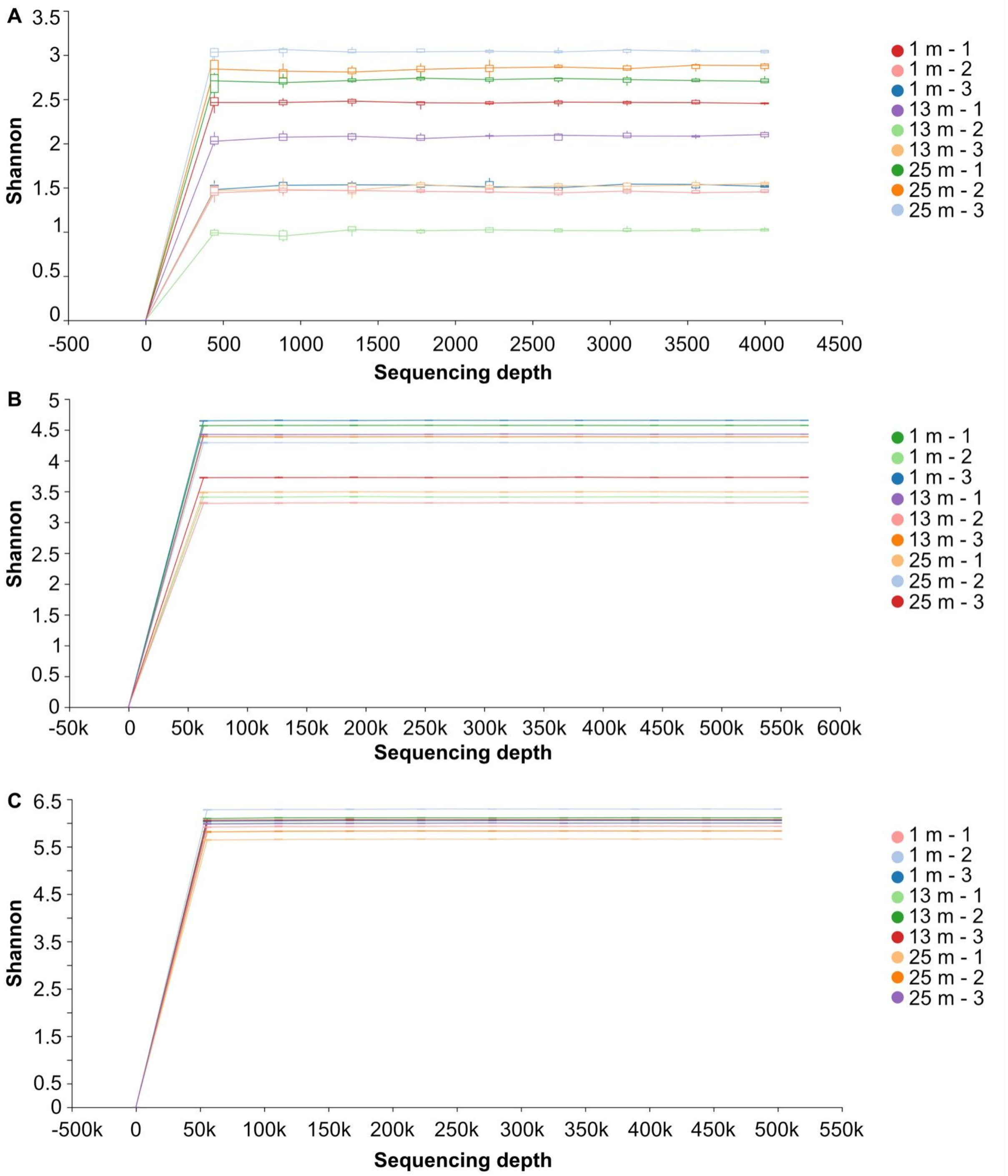


**Supplementary figure 1.** Alfa rarefaction per sample of the 12S (A), 16S (B) and COI (C) minibarcodes. Shannon index and the sequencing depth are represented by the y and x axes, respectively.

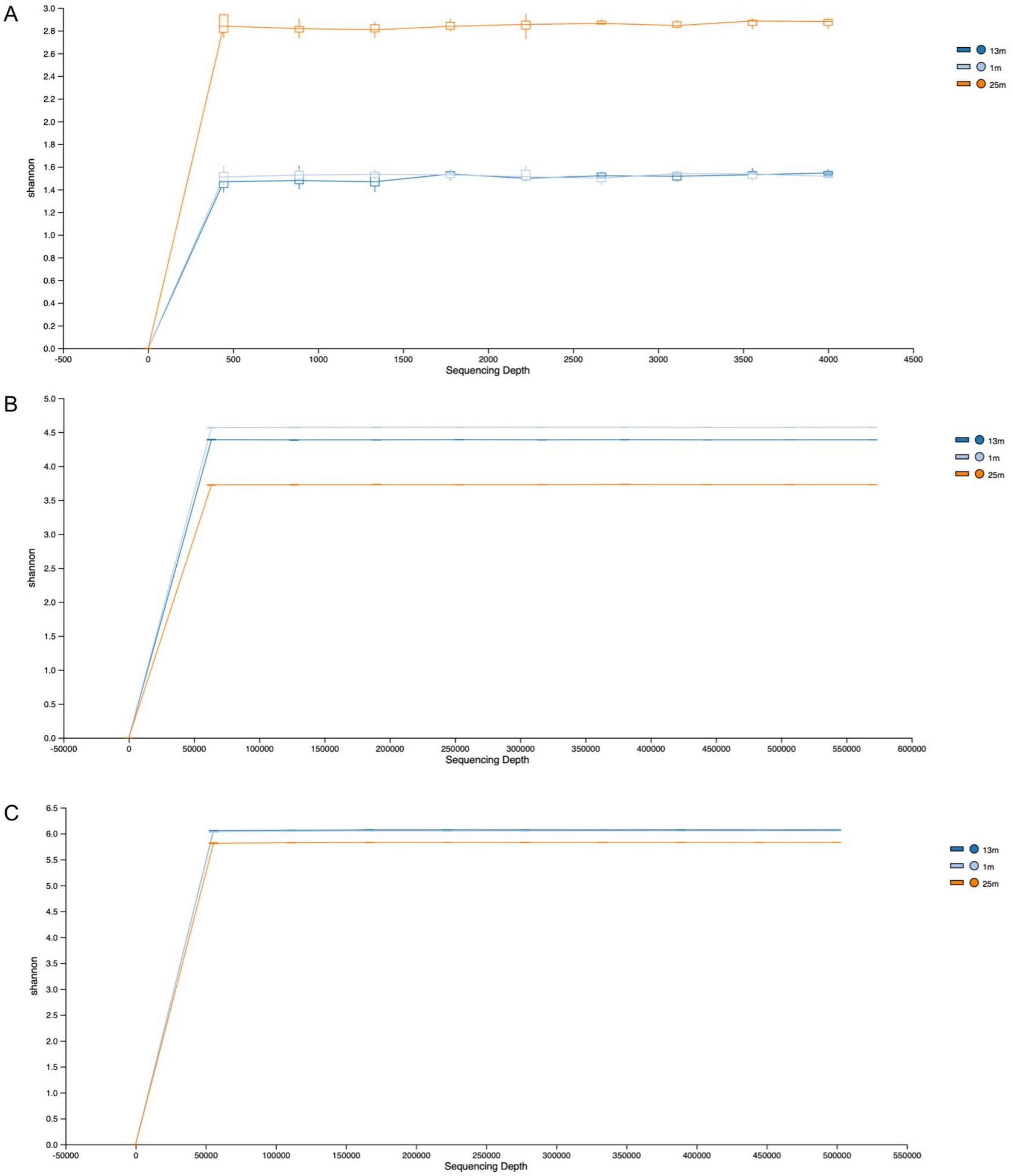


**Supplementary figure 2.** Alfa rarefaction by depth (1, 13 and 25 m) of the 12S (A), 16S (B) and COI (C) minibarcodes. Shannon index and the sequencing depth are represented by the y and x axes, respectively.

### 2. Fishes phylogenetic trees for each minibarcode

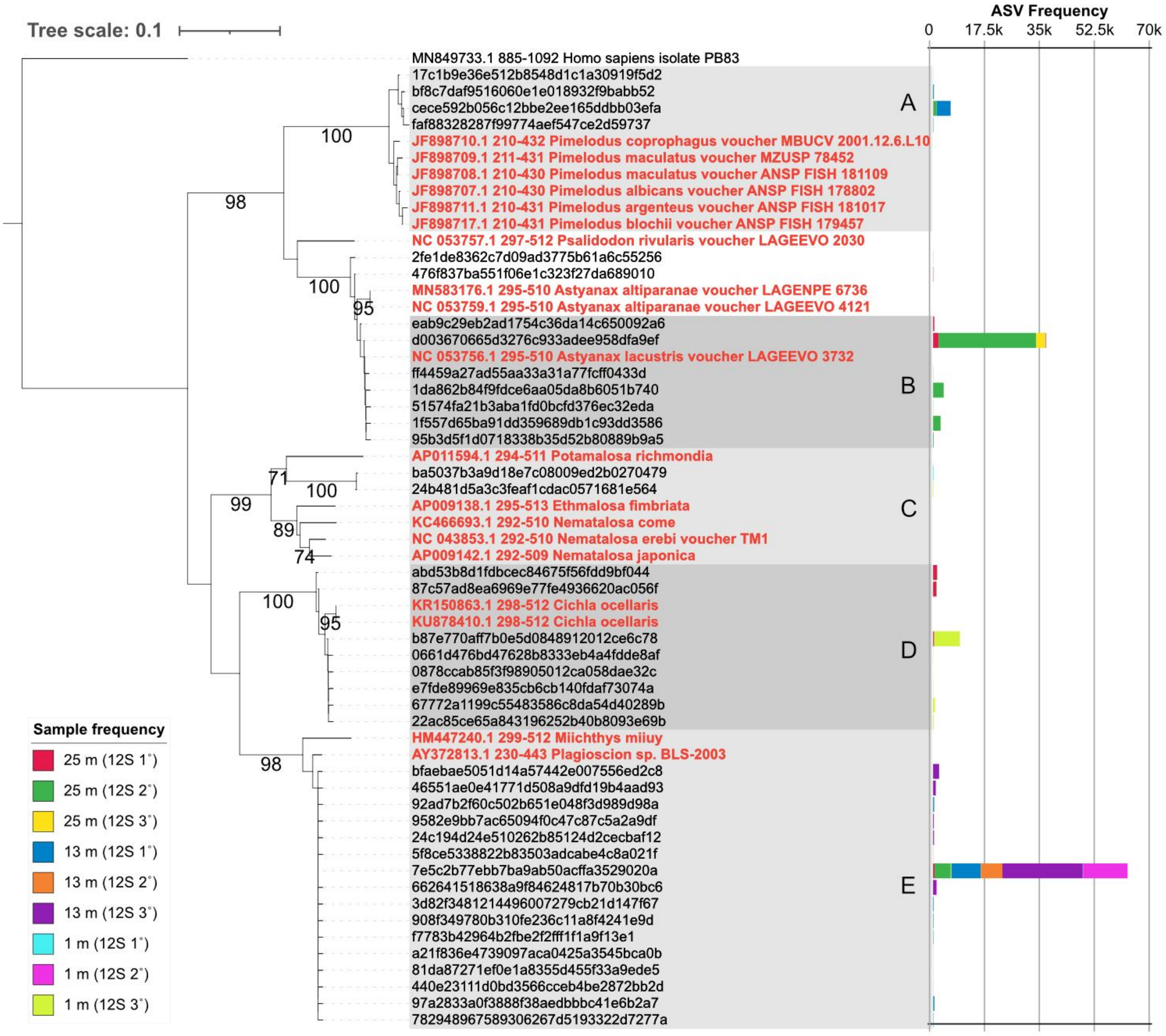


**Supplementary figure 3. Neighbor-joining tree using the minibarcode 12S region.** Leaves containing the ASVs belonging to fishes and reference sequences are coloured in black and red, respectively. *Pimelodus* sp. (Siluriforme) is represented by the clade A, *Astyanax lacustris* (clade B), Unknown clupeidae (Clade C), *Cichla ocellaris* (Clade D) and *Plagioscion* sp. (Clade E). Frequencies of each ASV within the replicates sampled at a depth of 1, 13 and 25 meters are shown at the right side of the leaves.

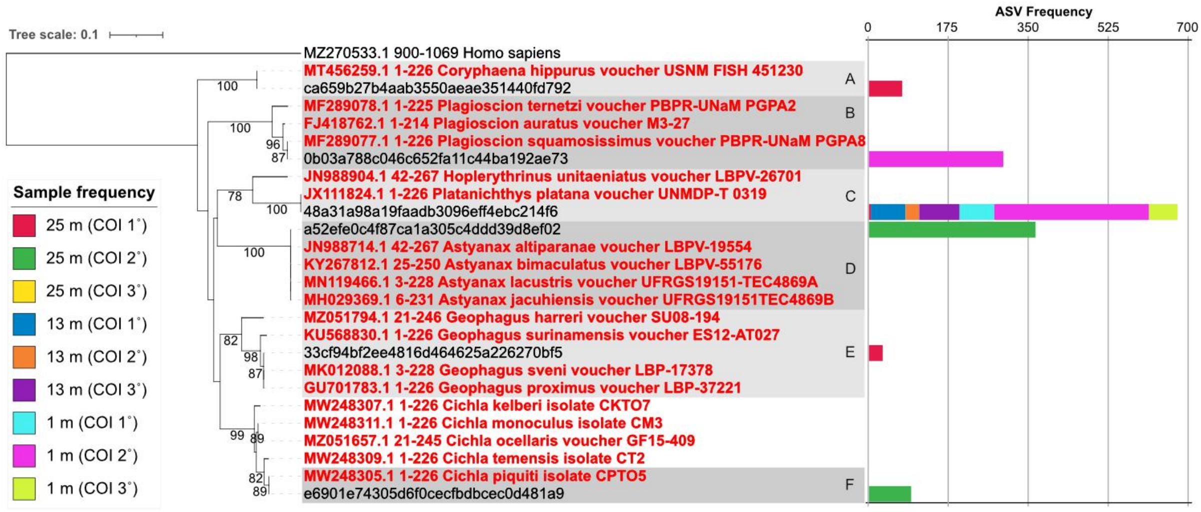


**Supplementary figure 4. Neighbor-joining tree using the minibarcode COI region.** Leaves containing the ASVs belonging to fishes and reference sequences are colored in black and red, respectively. *Coryphaena hippurus* is represented by the Clade A, *Plagioscion squamosissimus* (Clade B), *Platanichthys platana* (Clade C), *Astyanax* sp. (Clade D), *Geophagus* sp. (Clade E) and *Cichla piquiti* (Clade F). Frequencies of each ASV within the replicates sampled at a depth of 1, 13 and 25 meters are shown at the right side of the leaves.

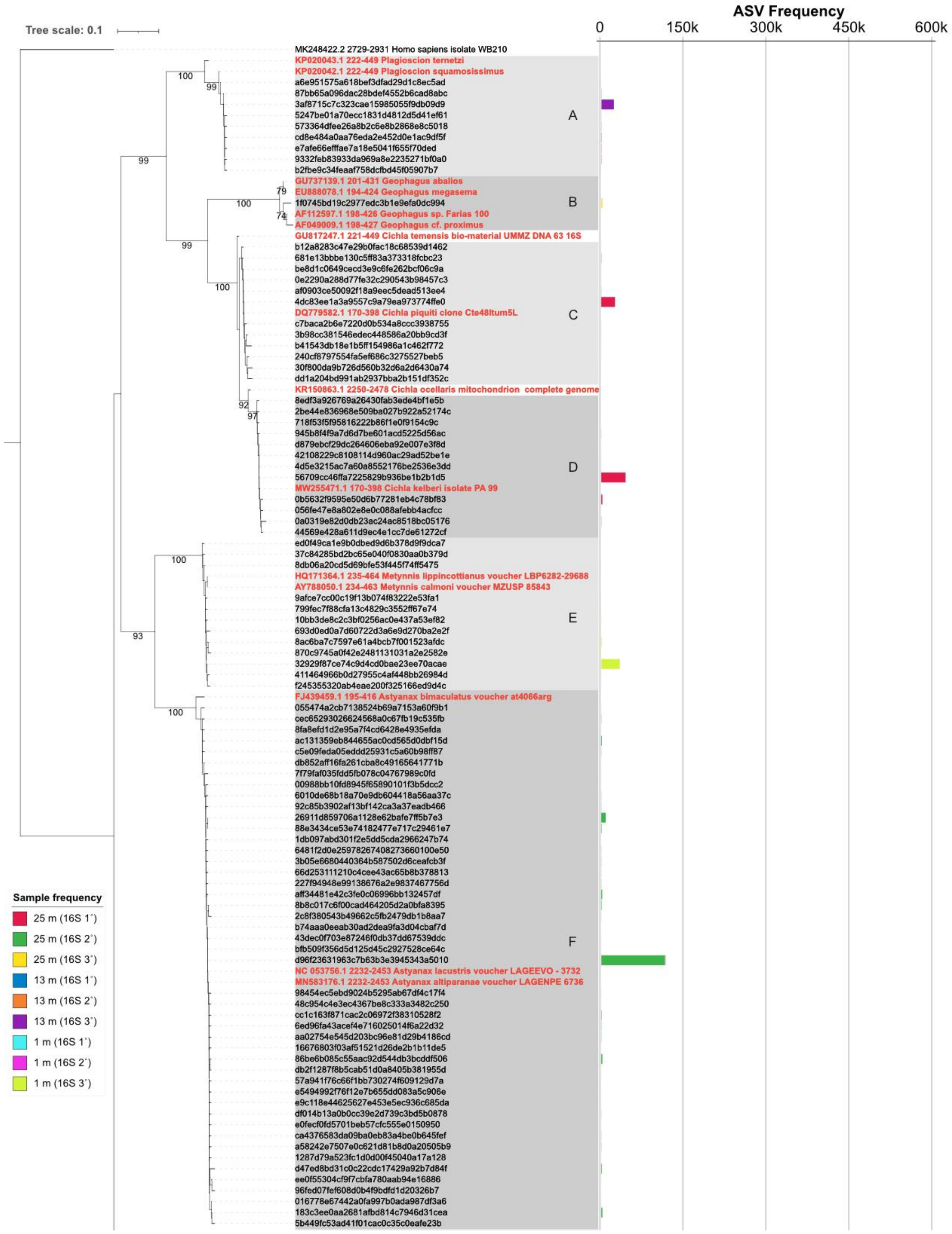


**Supplementary figure 5. Continued on the next page. Neighbor-joining tree using the minibarcode 16S region.** Leaves containing the ASVs belonging to fishes and reference species are coloured in black and red, respectively. *Plagioscion squamosissimus* is represented by the clade A, *Geophagus* sp. (Clade B), *Cichla piquiti* (Clade C), *Cichla kelberi* (Clade D), *Metynnis* sp. (Clade E) and *Astyanax* sp. (Clade F). Frequencies of each ASV within the replicates sampled at a depth of 1, 13 and 25 meters are shown at the right side of the leaves.

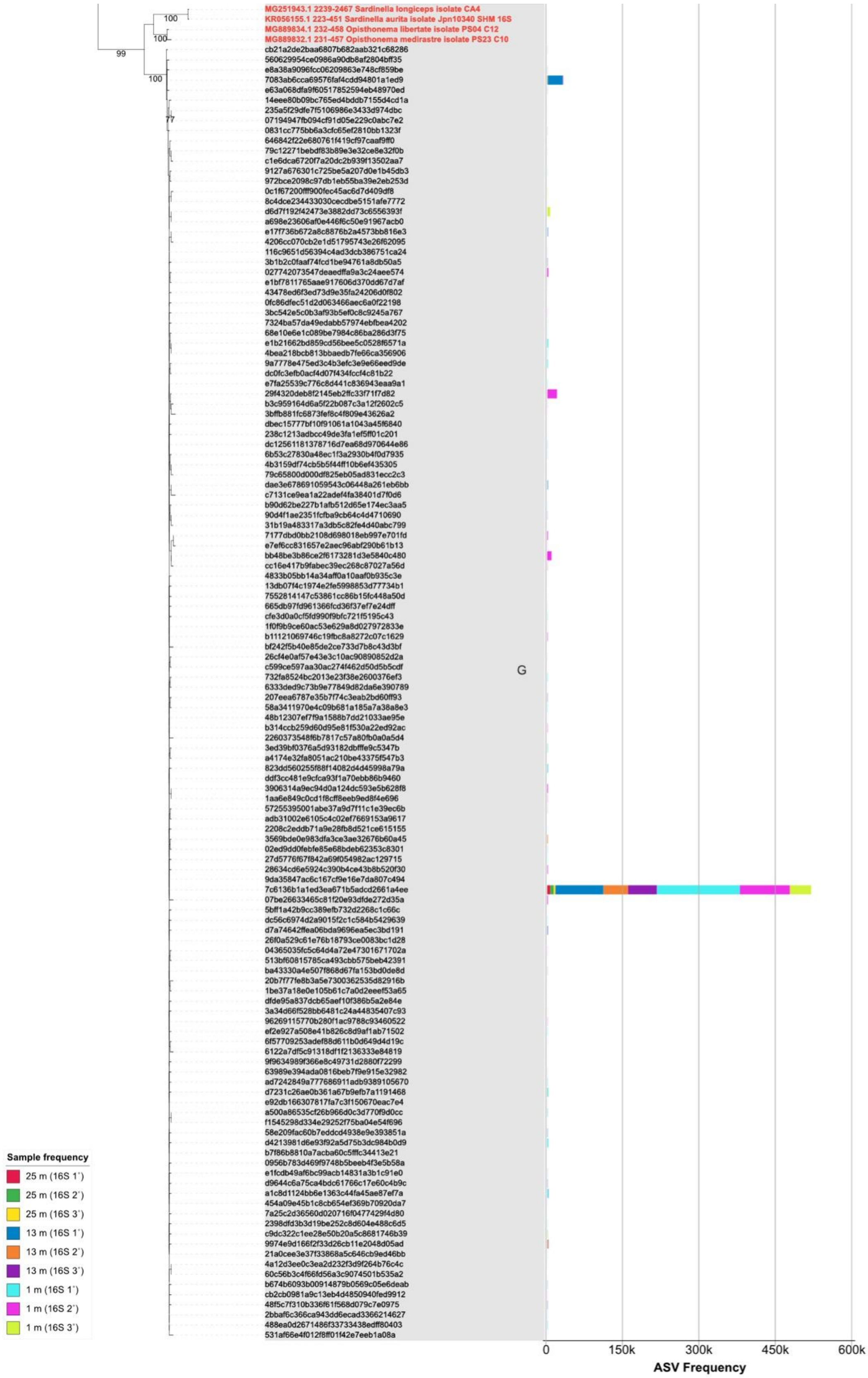


**Supplementary figure 5 (continued). Neighbor-joining tree using the minibarcode 16S region.** Leaves containing the ASVs belonging to fishes and reference species are colored in black and red, respectively. The frequency of each one within the triplicates sampled at a depth of 1, 13 and 25 meters is shown on the right side of the leaves.
