## Supplementary table 2 for "Environmental DNA from a small sample of reservoir water can tell volumes about its biodiversity"

**PRIMERS USED FOR DEEP SEQUENCING**

**Supplementary table 2**: Primer pairs used to amplify the target amplicons in the COI, 12S rRNA, and 16S rRNA genes from eDNA of the TI reservoir. Ta = Annealing temperature used.

| Target gene | Forward primer (5'-3') | Reverse Primer (5'-3') | Ta | Reference |
|---|---|---|---|---|
| COI | ACYAAICAYAAAGAYATIGGCAC | CTTATRTTRTTTATICGIGGRAAIGC | 48ºC | 3,4 |
| 12S rRNA gene | GTCGGTAAAACTCGTGCCAGC | CATAGTGGGGTATCTAATCCCAGTTTG | 57ºC | 5 |
| 16S rRNA gene | AYAAGACGAGAAGACCC | GATTGCGCTGTTATTCC | 53ºC | 6 |
